# Identification of activity-induced *Egr3*-dependent genes reveals genes associated with DNA damage response and schizophrenia

**DOI:** 10.1101/2020.09.01.276626

**Authors:** Ketan K Marballi, Khaled Alganem, Samuel J Brunwasser, Arhem Barkatullah, Kimberly T Meyers, Janet M Campbell, Robert R Mccullumsmith, Amelia L Gallitano

## Abstract

Bioinformatics and network studies have identified the immediate early gene transcription factor early growth response 3 (EGR3) as a master regulator of genes differentially expressed in the brains of patients with neuropsychiatric illnesses ranging from schizophrenia and bipolar disorder to Alzheimer’s disease. However, few studies have identified and validated *Egr3*-dependent genes in the mammalian brain. We have previously shown that *Egr3* is required for stress-responsive behavior, memory, and hippocampal long-term depression in mice. To identify *Egr3*-dependent genes that may regulate these processes, we conducted an expression microarray on hippocampi from wildtype (WT) and *Egr3-/-* mice following electroconvulsive seizure (ECS), a stimulus that induces maximal expression of immediate early genes including *Egr3*. We identified 71 genes that were differentially expressed between WT and *Egr3-/-* mice one hour following ECS. Bioinformatic analyses showed that many of these are altered in schizophrenia. Ingenuity pathway analysis revealed the GADD45 (growth arrest and DNA-damage-inducible) family (*Gadd45b, Gadd45g*) as the leading category of differentially expressed genes. Together with differentially expressed genes in the AP-1 transcription factor family genes (*Fos, Fosb*), and the centromere organization protein *Cenpa*, these results revealed that *Egr3* is required for activity-dependent expression of genes involved in the DNA damage response. Our findings implicate EGR3 as gene critical for the expression of genes that are disrupted in schizophrenia and reveal a novel requirement for EGR3 in the expression of genes involved in activity-induced DNA damage response.

---

Major advances in genetics and genomics over the past decade have led to identification of hundreds of genes associated with risk for neuropsychiatric illnesses such as schizophrenia, bipolar disorder, depression, and Alzheimer’s disease. Many of these risk genes are shared across these disorders ^1^, each of which is characterized by cognitive dysfunction. With the discovery of such a vast number of putative illness-influencing genes, the challenge becomes how to identify the functional relationships among them, which should provide insight into the mechanisms underlying neuropsychiatric and neurodegenerative illnesses.

A promising approach has been to define functional networks of genes that are differentially regulated in individuals affected by these illnesses, compared with controls, and then to identify the “Master Regulatory Genes” that best account for these differences in gene expression. The immediate early gene transcription factor early growth response 3 (EGR3) has emerged as such a master regulator of differentially expressed genes (DEGs) in multiple neuropsychiatric disorders including schizophrenia ^2^, bipolar disorder ^3^, and most recently Alzheimer’s disease ^4^. However, despite these regulatory relationships identified using bioinformatic approaches, few genes regulated by EGR3 have been validated in the brain *in vivo*.

One of the first EGR3 downstream target genes to be identified in the brain is activity-regulated cytoskeleton-associated protein (*Arc*) ^5^. ARC has been implicated in risk for schizophrenia by studies of rare variants, de novo mutations, and single nucleotide polymorphism association ^6-9^. EGR3 is also reported to upregulate glutamic acid decarboxylase A4 (GABRA4) in response to seizure ^10, 11^. Neuropathologic studies have identified dysfunction in the GABAergic system in schizophrenia and GABRA4 is an autism susceptibility gene ^12^.

Our prior work has identified deficits in the function of N-methyl D-aspartate (NMDA) receptors in *Egr3*-/- mice, specifically those containing the NR2B subunit ^13^. This indicates that *Egr3* is required for function of a receptor at the center of one of the leading models of schizophrenia pathogenesis, the NMDA receptor hypofunction model of schizophrenia ^14^. In this and other studies we found that *Egr3*-/- mice have deficits in stress-responsive behavior, memory, and hippocampal long-term depression, further supporting the importance of *Egr3* in behavioral and electrophysiologic processes implicated in neuropsychiatric disorders and cognitive processes ^13, 15^.

Based on these findings, we hypothesized that EGR3 is a critical transcriptional regulator in a biological pathway of proteins essential for memory, synaptic plasticity, and the risk for schizophrenia ^16^. As an immediate early gene EGR3 expression is induced in response to neuronal activity in a manner dependent upon NMDA receptor function and calcium signaling, processes implicated in schizophrenia ^17^. EGR3 interacts in regulatory feedback loops with other EGR-family genes in the immune system, including EGR1, EGR4 and NAB2, each of which maps to GWAS loci for schizophrenia ^18-20^. And dysfunction in these genes leads to abnormalities in processes that are disrupted in schizophrenia, including memory, synaptic plasticity, immune function, growth factor-mediated processes, myelination, and vascularization ^13, 16,21-28^. Based on the central role of EGR3 in these critical processes, we hypothesized that genes regulated downstream of EGR3 will contribute to risk for schizophrenia and other neuropsychiatric disorders that are characterized by abnormalities in cognition, memory, and synaptic function.

To test this hypothesis, we sought to characterize the complement of genes that require EGR3 in the hippocampus, a critical region for memory formation. We used electroconvulsive seizure (ECS) to maximally activate immediate early gene expression in the hippocampus of *Egr3-/-* and wildtype (WT) mice and conducted an expression microarray to identify genes differentially expressed between the genotypes one hour and two hours following the stimulus, compared to baseline unstimulated conditions. Here we show that over 71 genes are differentially expressed in the hippocampi of *Eg3*- /- mice compared to WT controls. Numerous of these *Egr3*-dependent genes map to schizophrenia GWAS loci, and are abnormally expressed in the brains of patients with schizophrenia, bipolar disorder, depression, and Alzheimer’s disease, supporting findings of studies indicating that EGR3 may be a master regulator of genes involved in risk for numerous neuropsychiatric disorders ^2-4^. This approach has also revealed the novel finding that one of the major classes of genes regulated downstream of EGR3 in the hippocampus are genes involved in the DNA damage response.

## Materials and Methods

### Mice

Previously generated *Egr3-/-* mice ^29^ were backcrossed to C57BL/6 mice for greater than 20 generations. All studies were carried out on homozygous adult progeny (*Egr3*-/- and wildtype (WT)) resulting from heterozygote matings and were assigned as “matched pairs” at the time of weaning. Matched pairs were subjected to identical conditions for all studies. The microarray studies, and quantitative RT-PCR validation studies, were performed on a cohort of male mice ages 6-12 months (n = 4 per group). Replication studies were performed on a cohort of female mice, ages 12 – 15 months (n = 4 – 5 per group). Animals were housed on a 12-hour light/dark cycle with *ad libitum* access to food and water. All studies were performed in accordance with the University of Arizona, Institutional Animal Care and Use Committee (IACUC). This study was carried out in accordance with the recommendations of IACUC guidelines, IACUC.

### Electroconvulsive Seizure and Tissue Collection

Electroconvulsive stimulation was delivered to mice via corneal electrodes 5 min.s following application of 0.5% proparacaine hydrochloride ophthalmic solution (Akorn, Inc., Lake Forest, IL, United States). The cohort of mice used for the microarray study underwent ECS without general anesthesia. The replication cohort of mice underwent ECS following general anesthesia. Isoflurane anesthesia (VetOne, Boise, ID, United States) was administered in an enclosed chamber at a flow rate of 0.5 mL/min in oxygen. Animals were removed from the chamber after 2 minutes of complete anesthetization, transferred to room air to recover to a level of light anesthesia, and then administered electrical stimulation of 260 A for 1 ms duration and a pulse width of 0.3 mm, 1 ms (Ugo Basile, Varese, Italy) via orbital electrodes. Mice were observed to undergo tonic-clonic seizure and were placed in their home cage to recover for one hour prior to sacrifice. Control animals remained in their home cages undisturbed until the time of sacrifice.

### Tissue Collection and RNA Isolation

Animals were sacrificed using isoflurane overdose, followed by decapitation. The brains were removed, rinsed in ice-cold phosphate buffered saline (PBS), and hemisected along the central sulcus into right and left hemispheres for further studies to quantify mRNA expression. Whole hippocampi were rapidly dissected and immediately placed in RNAlater (Ambion, Waltham, MA, United States). Tissue was transferred to 1.5-mL Eppendorf tubes, frozen on dry ice and then stored at -80°C. For the microarray and follow-up qRT-PCR studies, RNA was isolated using TRIzol reagent (Life Technologies, Carlsbad, CA, United States) per the manufacturer’s protocol. RNA was resuspended in RNAse-free water and quantitated by spectrophotometry. RNA quality and concentration were determined using an Agilent Bioanalyzer 2100 prior to microarray analysis and reverse transcription for qRT-PCR. An aliquot of the RNA samples was sent to the Microarray Resource Center, Yale/NIH Neuroscience Microarray Center (New Haven, CT, United States) for analysis using an Illumina Mouse WG6 v3.0 expression beadchip microarray. For the replication cohort, RNA isolation was performed using TRI reagent (Sigma-Aldrich, St. Louis, MO, United States) and MagMaxTM Total RNA isolation kit (Ambion, Waltham, MA, United States) according to the manufacturer’s protocol, and quantified using the NanoDrop ND-1000 spectrophotometer (Thermo Scientific, Waltham, MA, United States).

### Microarray Procedure and Analysis

Gene expression analysis was performed using Illumina Mouse WG6 v3.0 expression beadchip microarray and analyzed using two independent microarray analysis methods. Data analysis and quality control was initially performed using Gene Pattern (Reich et al., 2006), with normalization using the cubic spline method under the following settings (FDR < 0.05) to determine significantly differentially expressed genes between the WT and *Egr3*-/- groups 1 hour following ECS. A parallel analysis was performed using the Illumina Genome studio 2010 software to identify DEGs using the following settings: background subtraction, quantile normalization, *p* < 0.05, Illumina custom algorithm. The Illumina custom algorithm uses “Diff scores” to account for reproducibility of results, by P value and the magnitude of gene expression difference represented by signal intensity between the reference and control groups ^30^, where Diff score = 10 X (*Egr3-/-* ECS signal intensity gene A - WT ECS signal intensity gene A) x log10 P value, where a P value > 0.05, would correspond to a diff score of > 13 and < -13. Finally, a list of common DEGs between both programs was generated. From this common DEG list, genes that showed a fold change difference of 1.4-fold or higher between the two groups and a p value of < 0.05 (Table 1), were used for all subsequent analyses ^31^.

**Table 1.**
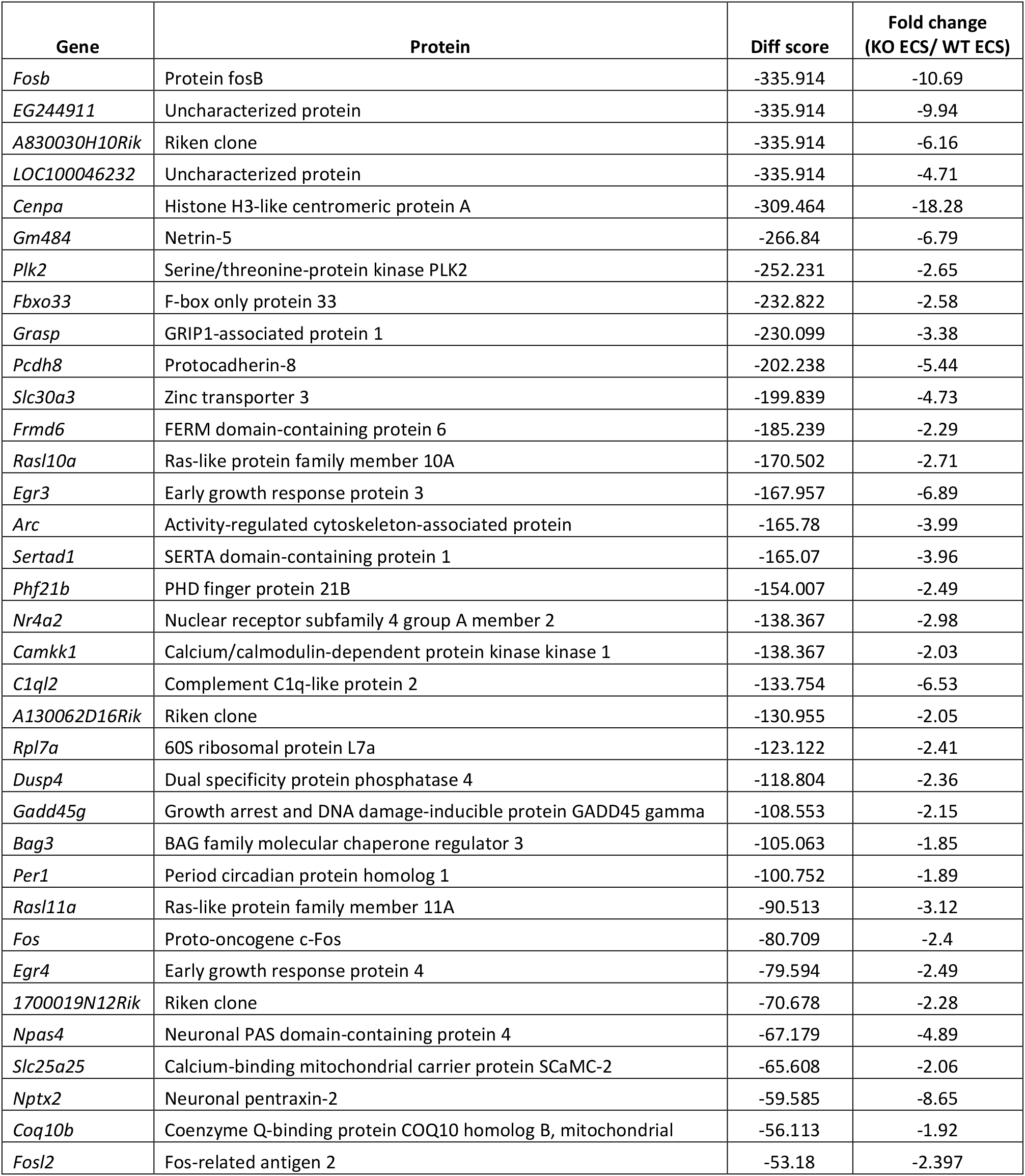
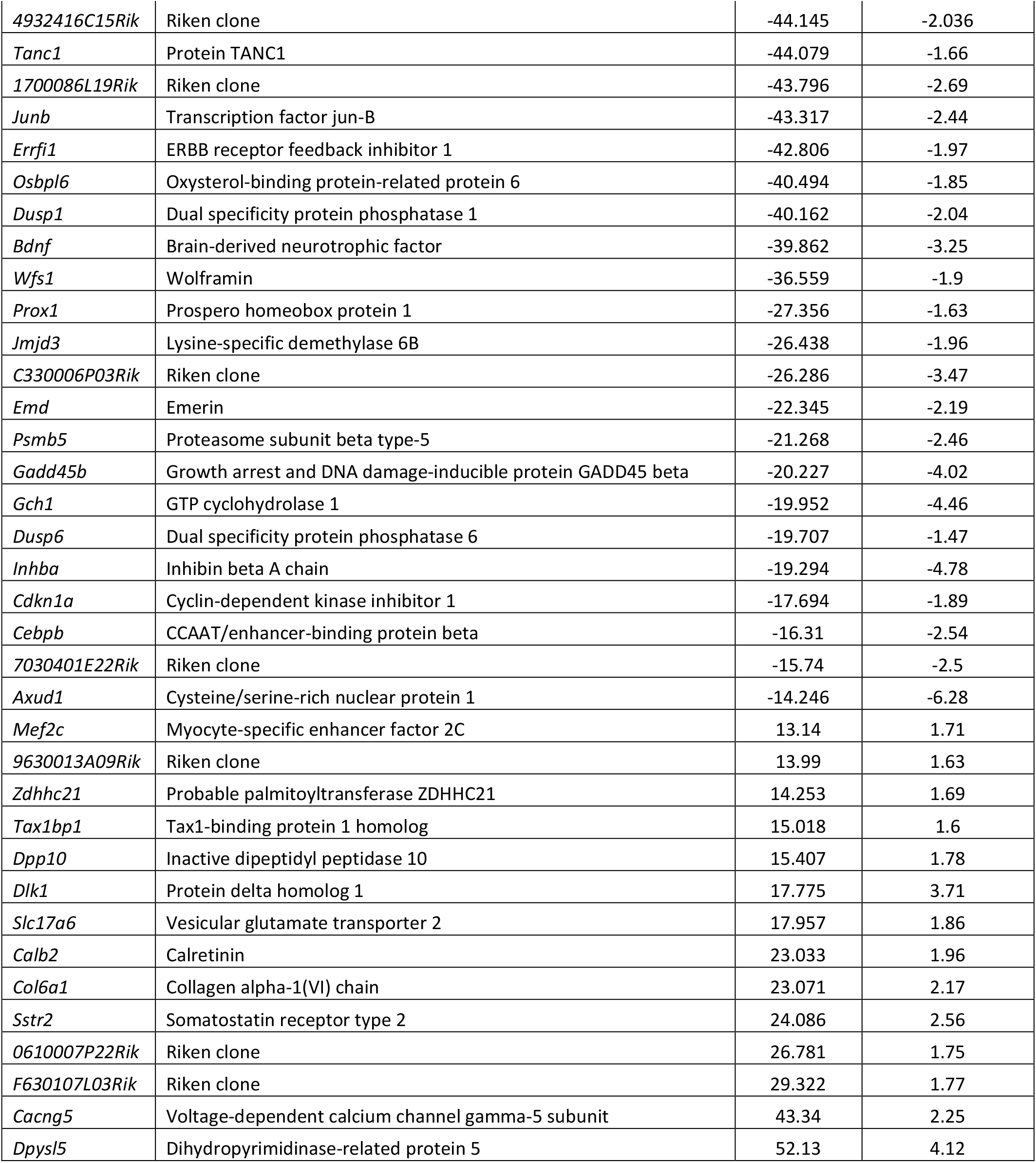
Genes differentially expressed in the hippocampus of WT versus *Egr3*-/- mice following ECS. 71 genes showed a fold change difference of ≥1.4 fold between the two groups with a p value of < 0.05.

### Ingenuity Pathway Analysis

The 71 DEGs list was imported into IPA (Ingenuity Systems, Redwood City, CA, USA; www.ingenuity.com), and analyses were carried out using default settings with the microarray WG6 gene set as background. IPA utilizes a right tailed Fisher’s exact test to calculate p values that are reflective of the possibility that overlap between input data and a given process or pathway, generated by curated data from the IPA database, is nonrandom.

### qRT-PCR

For qRT-PCR studies, mRNA was reverse transcribed into cDNA, as previously described ^32^, and used as a template for qRT-PCR using FastStart SYBR Green Master mix (Roche Applied Science, Indianapolis, IN, United States) on a 7500 Fast Real-Time PCR machine (ThermoFisher Scientific, Waltham, MA, United States). Each sample was amplified in triplicate for the gene of interest and the housekeeping gene phosphoglycerate kinase 1 (*Pgk1*). *Pgk1* was selected as a housekeeping gene as it showed no significant changes in gene expression across experimental groups in the microarray data. This was validated by qRT-PCR across both male and female cohorts. Fold changes in gene expression were calculated and data were plotted using the 2^-ΔCT^method ^33^.

### Statistical analysis

Details of microarray and IPA analysis statistics are described in the respective sections. For analysis of qRT-PCR data, we utilized a two-way analysis of variance (ANOVA) followed by Tukey’s *post hoc* test using GraphPad Prism with a significance threshold of p < 0.05.

## Results

To identify candidate target genes of EGR3 we conducted an expression microarray of genes differentially expressed in the hippocampus of *Egr3*-/- mice compared with WT controls. Because EGR3 is an activity dependent transcription factor, we used electroconvulsive stimulation to induce a seizure, which maximally actives expression of immediate early genes in the brain ^17^. Hippocampal RNA was isolated from *Egr3*-/- and WT mice under three conditions: baseline (no ECS), one hour after ECS, and two hours after ECS. This allowed us to identify genes that require *Egr3* under basal conditions as well as following neuronal activity.

Figure 1A shows a heatmap of the 71 genes that are differentially expressed between *Egr3*-/- and WT mice one hour following ECS, the timepoint with the maximum number of DEGs. Of these, 57 genes were downregulated in *Egr3-/-* mice, while 14 genes were upregulated in *Egr3-/-* mice, compared to WT mice. Table 1 lists the relative expression levels for each DEG identified at 1 hr. after ECS.

**Figure 1.**
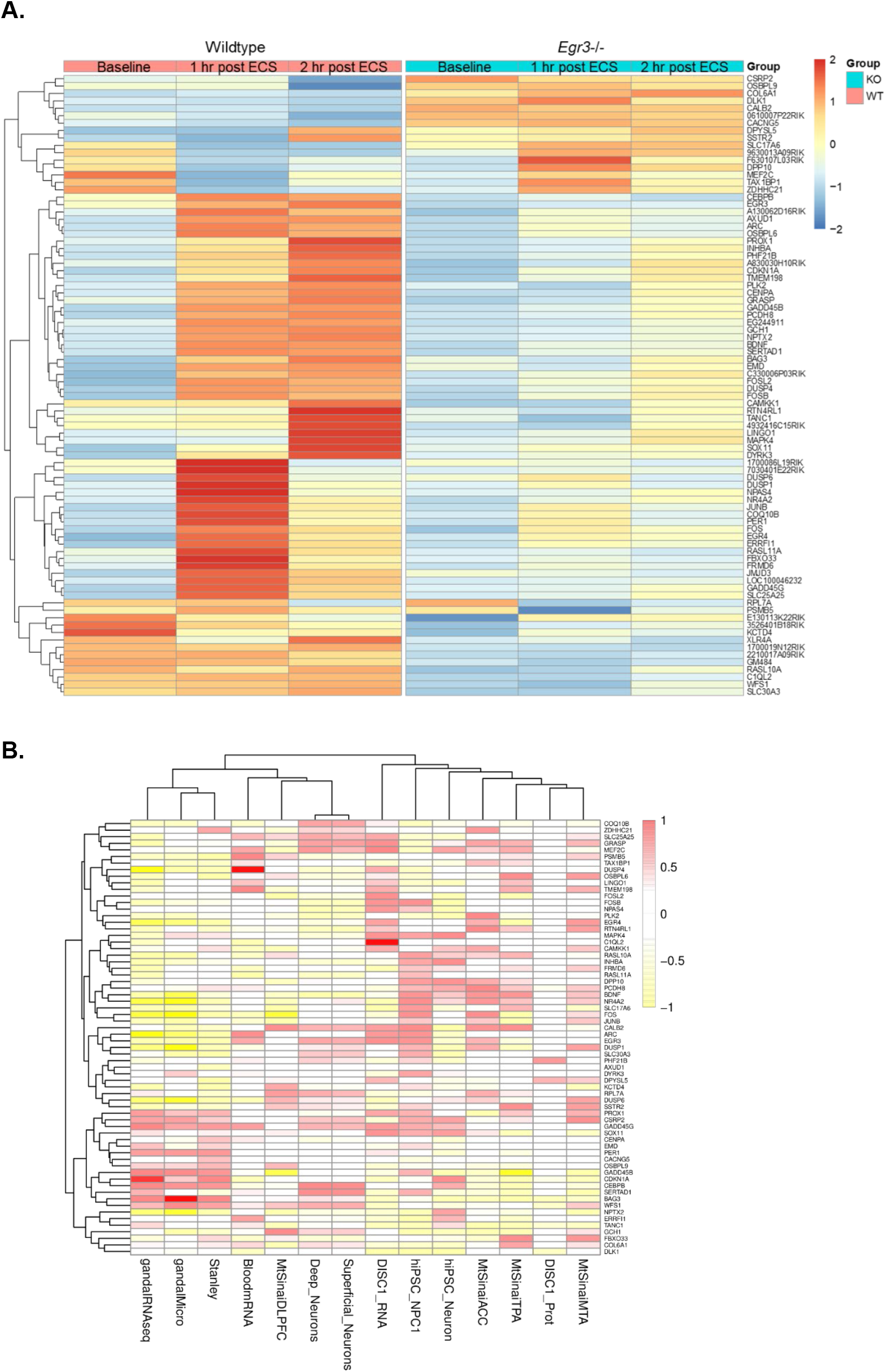
EGR3-dependent genes show altered expression in schizophrenia studies. **A**. Expression heatmap of genes differentially expressed in WT versus *Egr3*-/- mice following ECS. Expression microarray revealed 71 genes that were differentially expressed in the hippocampus of *Egr3*-/- mice compared to WT one hour following ECS. The average expression level for each of these genes is shown along a horizontal row in WT (left) and *Egr3*-/- mice (right) at baseline, and 1 hr and 2 hrs following ECS. (n = 4 animals per condition). **B**. Fourteen published gene expression studies in schizophrenia were queried for the 71 genes that are differentially expressed in *Egr3*-/- compared with WT mouse hippocampus 1 hr following ECS. The heatmap shows normalized log2 fold change values (schizophrenia vs. control) from the 14 published study datasets (vertical columns) for each of the 71 EGR3-dependent genes (horizontal rows).

The use of three timepoints (baseline, and 1 and 2 hours after ECS) allowed us to perform cluster analyses of 84 DEGs. Supplemental Figure S1 shows four major clusters of DEGs based on the patterns of gene expression changes in WT versus *Egr3*-/- mice over time.

### *Egr3*-dependent genes are altered in schizophrenia studies

Next, we tested the original hypothesis for our study, that EGR3 regulates genes that play a role a role in schizophrenia. To do this we examined the expression levels of each of the 71 genes that were differentially expressed at 1 hour following ECS in 14 published schizophrenia study datasets. These include studies of genes differentially expressed in post-mortem brains, peripheral blood, fibroblasts, and induced pluripotent stem cells, from schizophrenia patients compared with controls. Expression levels for each of the 71 DEGs were assessed in the data from each schizophrenia study and used to create a heatmap for *in vivo* expression (Table 2, Fig. 1B).

**Table 2.**
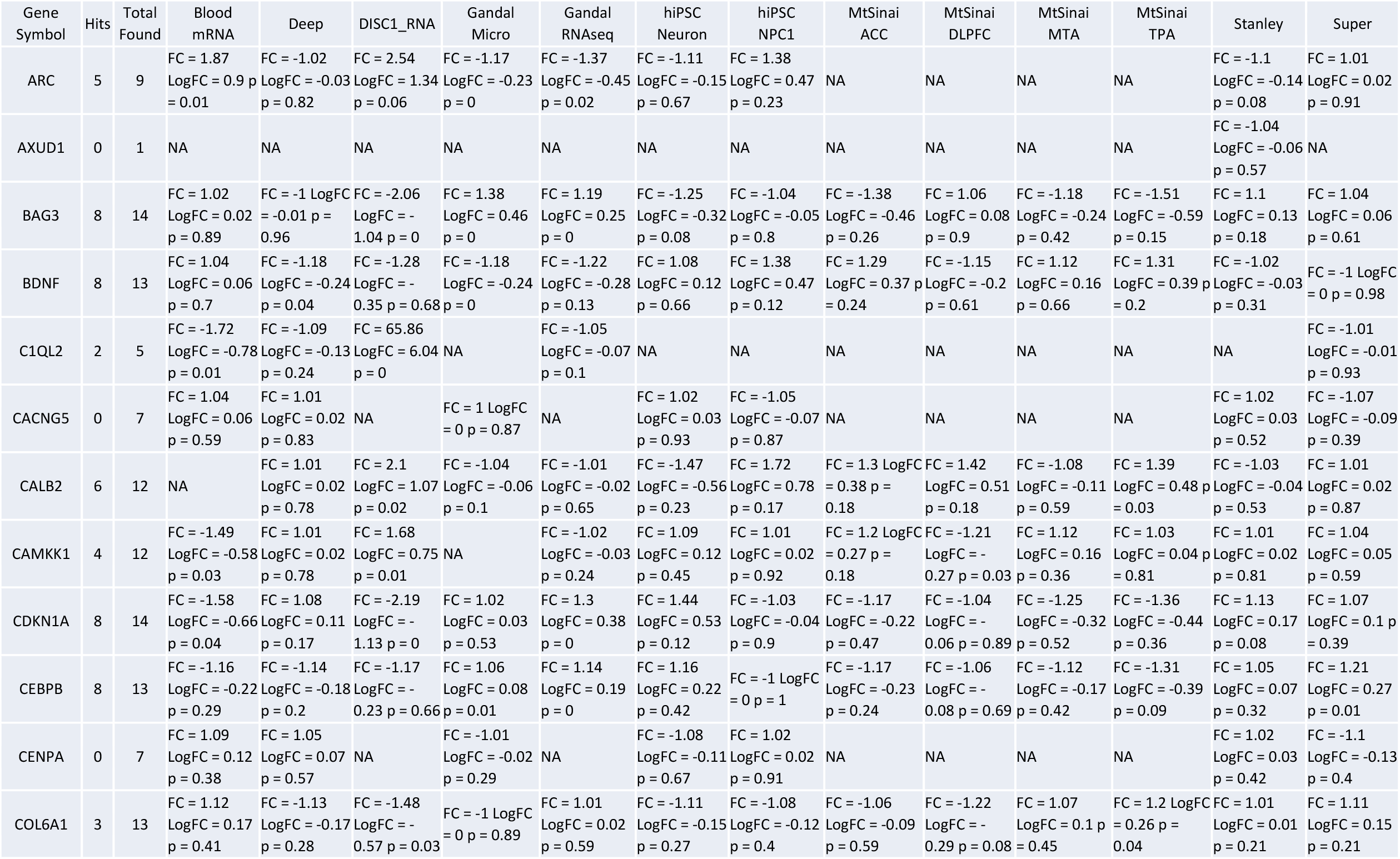
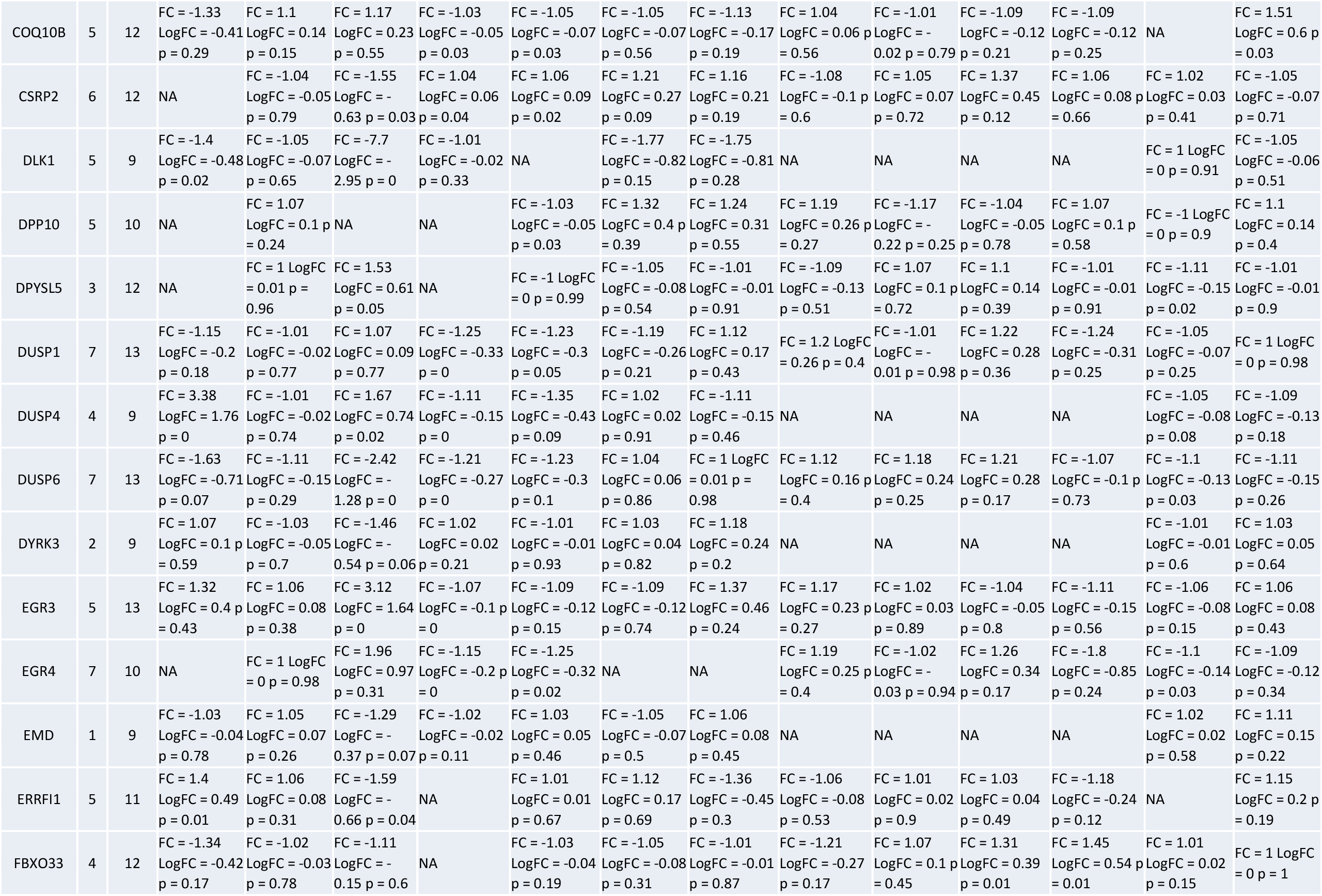
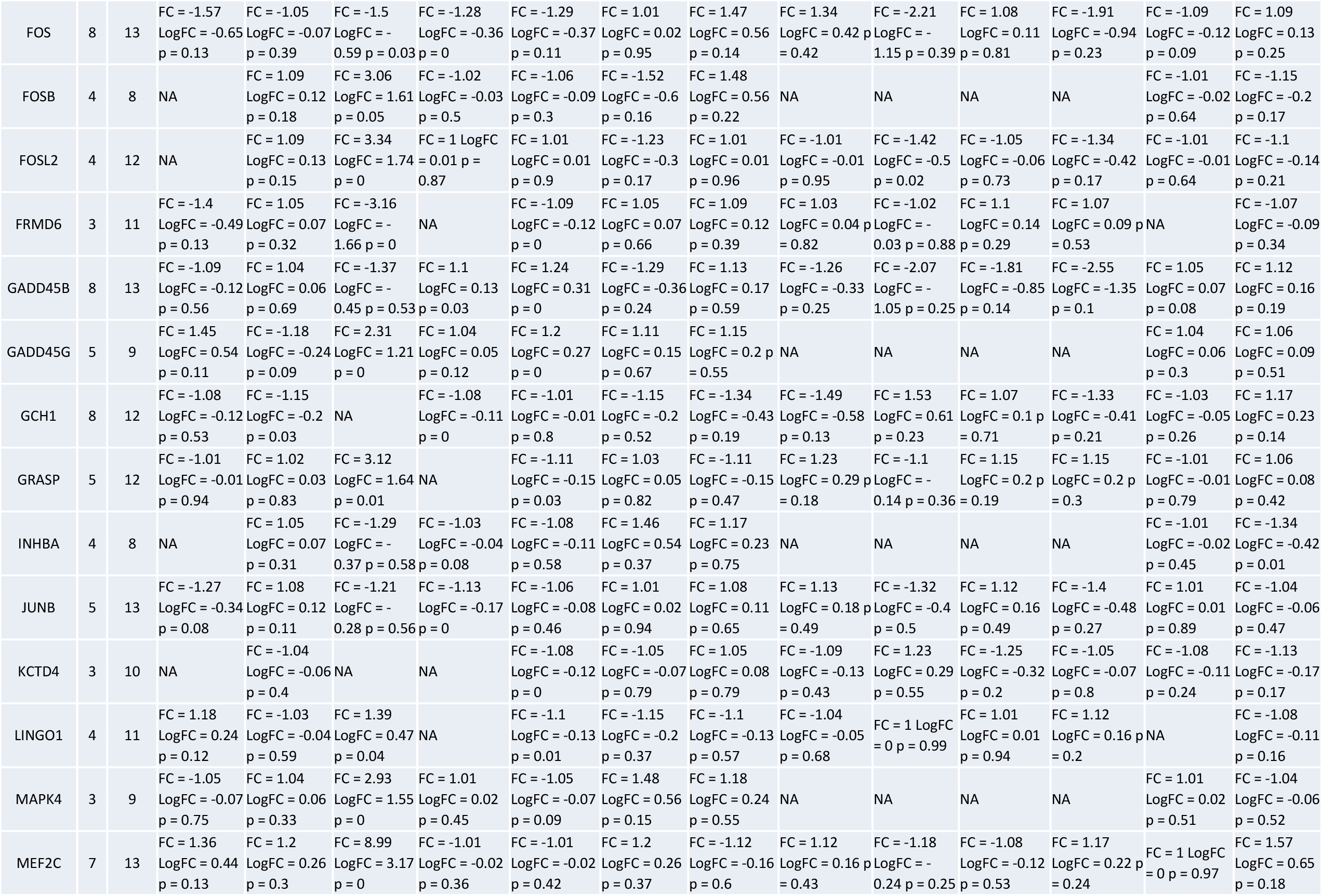
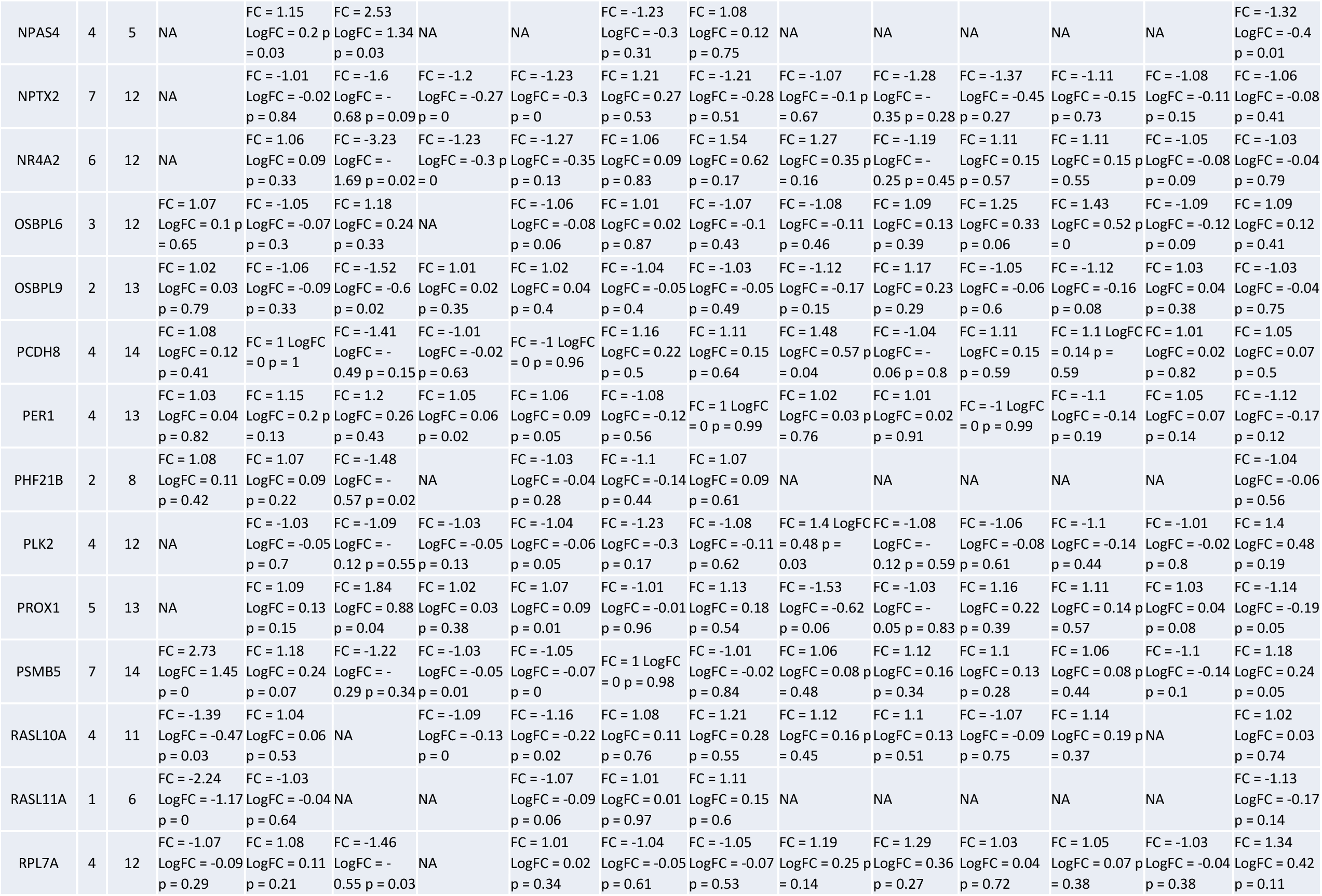
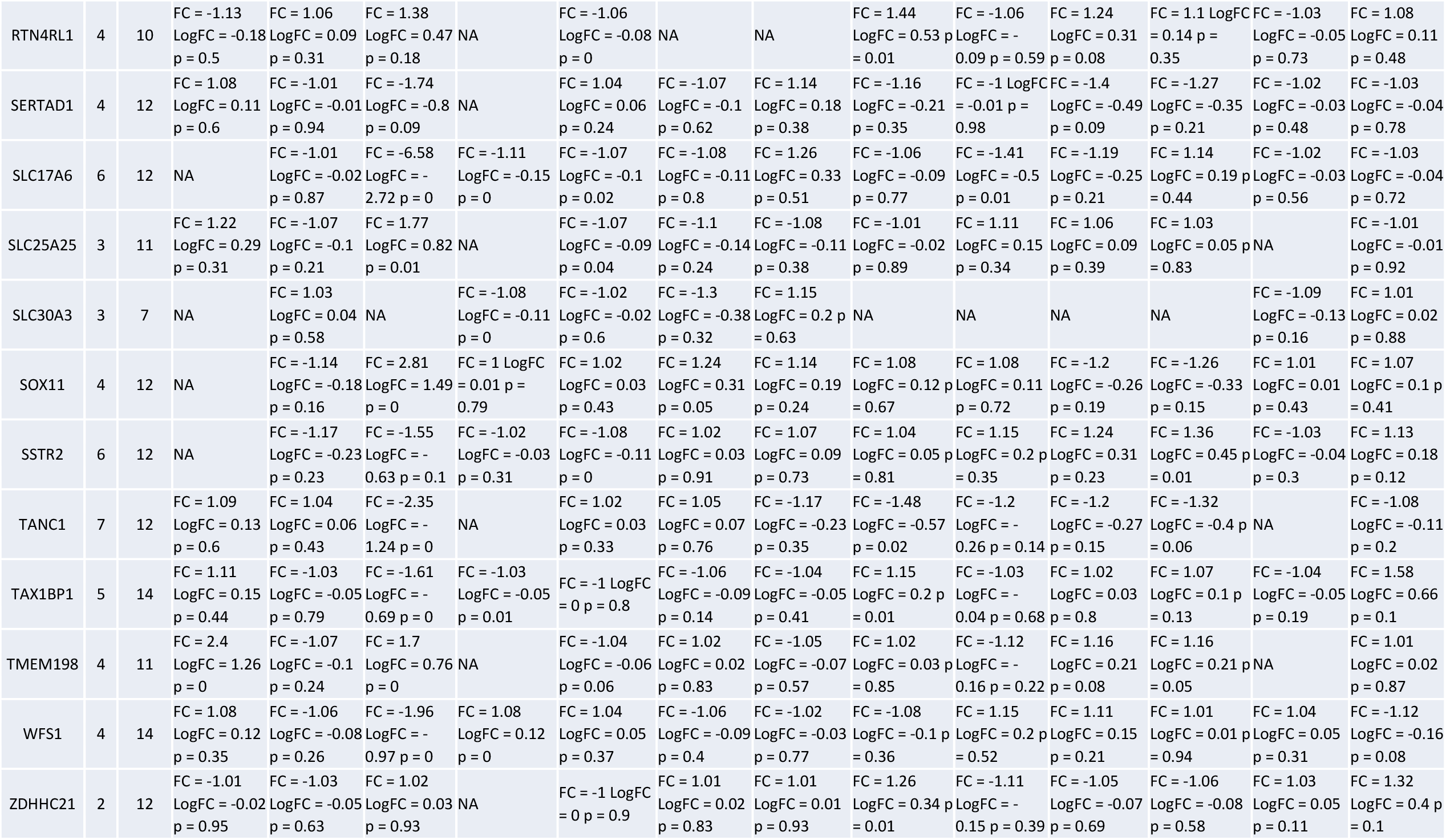
EGR3-dependent differentially expressed genes show altered expression in schizophrenia studies.

Supplemental Figure S2 shows the proportion of schizophrenia datasets in which each of the 71 DEGs was identified as significantly different between schizophrenia subjects and controls. All but three genes coincided with differentially expressed genes in schizophrenia. These findings support the hypothesis that EGR3 regulates genes that are abnormally expressed in schizophrenia.

### EGR3-dependent DEGs are involved in DNA damage response and behavior

To identify the major biological pathways regulated by EGR3 in the hippocampus we conducted a canonical pathway analysis using the Ingenuity Pathway Analysis (IPA) program. The results showed that the most significantly overrepresented pathway in the DEG list was GADD45 (Growth Arrest and DNA Damage) signaling, followed by corticotropin signaling, p53, ATM and Jak/STAT signaling, respectively (Fig. 2A). A literature survey revealed that GADD45, p53, ATM and Jak/STAT pathways are all involved in DNA damage response ^34 35, 36^. In addition, we also observed that 26 genes in the DEG list have roles in the regulation of behavior (Fig. 2B). Based on these observations we chose to follow-up the microarray results with validation of genes that were relevant to DNA damage response and/or neuronal function in the context of behavior.

**Figure 2.**
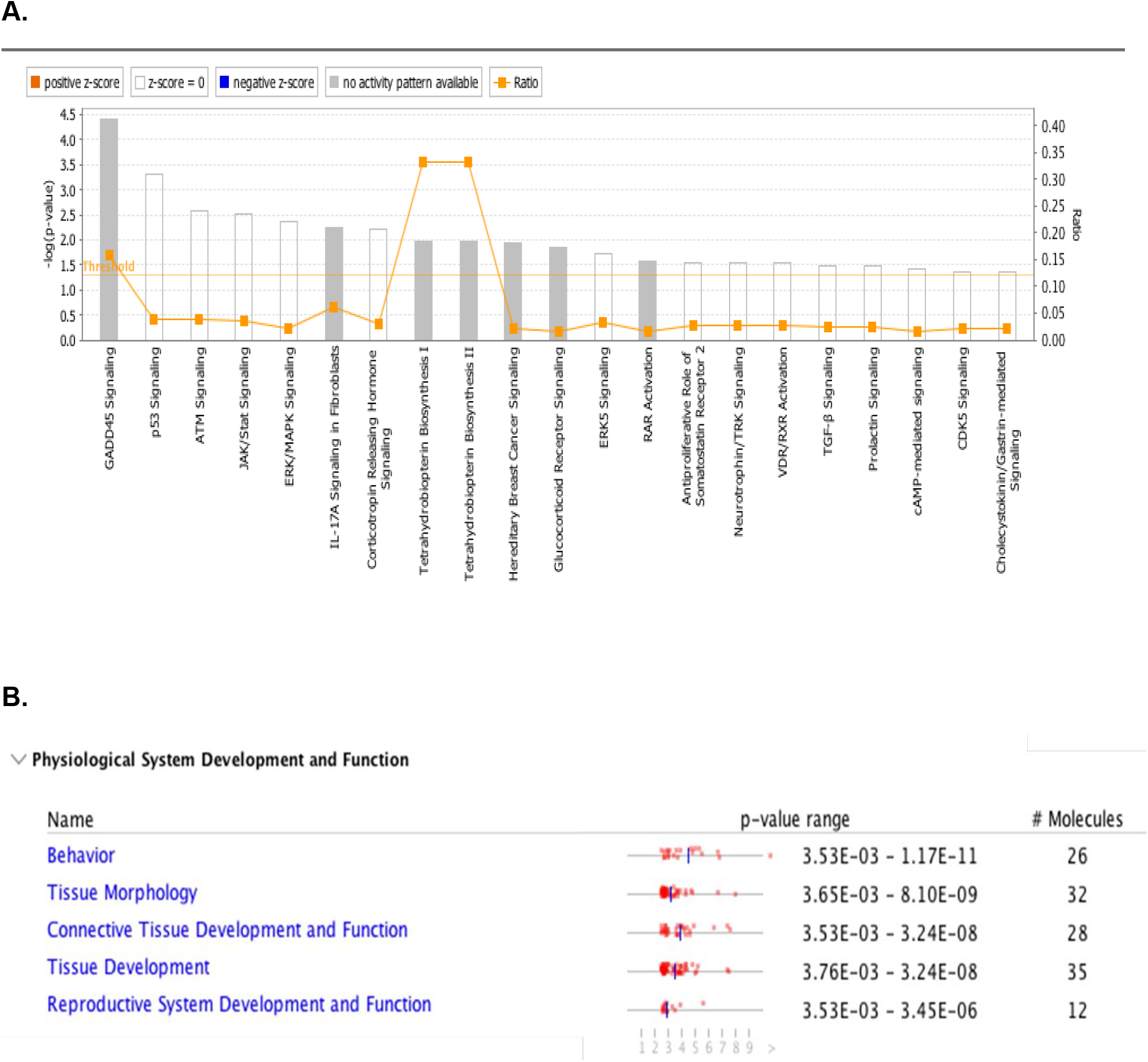
Top canonical pathways and physiological function categories associated with DEGs generated by IPA. Ingenuity Pathway “core analysis” performed on microarray results reveals pathways associated with the genes that are differentially expressed in hippocampus from *Egr3*-/- vs. WT mice following ECS. **A**. “Threshold” indicates the minimum significance level [scored as –log (p-value) from Fisher’s exact test, set here to p < 0.05]. “Ratio” (yellow points) indicate the number of molecules from the DEG dataset that map to the pathway listed, divided by the total number of molecules that map to the canonical pathway from the IPA knowledgebase. **B**. DEGs are most significantly enriched for genes involved in regulation of behavior. The top 5 scoring pathways in Physiological System Development and Function are shown with significance ranges for subcategories within each hit (p-value range) and number of molecules associated.

### GADD45 signaling genes require *Egr3* for ECS-induced expression

The GADD45 family consists of proteins involved in regulation of DNA repair ^37, 38^ (Fig 3A), DNA demethylation ^39^, neurogenesis ^40^ and response to stress ^41^. Since GADD45 signaling was significantly overrepresented in the DEG list, we conducted follow up studies on the members of the GADD45 signaling pathway that were in our DEG list: *Gadd45b* and *Gadd45g*.

**Figure 3.**
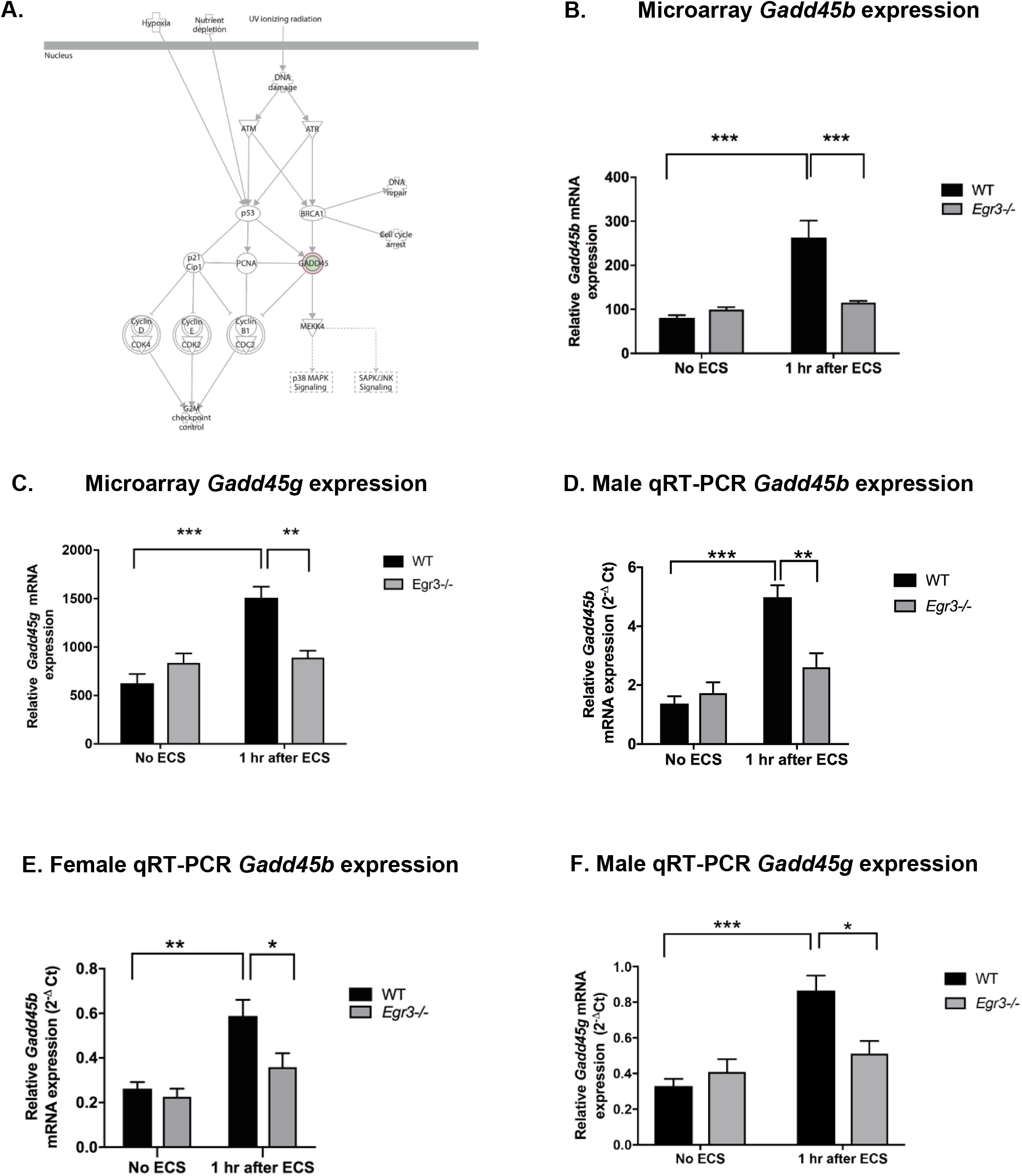
GADD45 family genes are differentially expressed in *Egr3*-/- mice. A. GADD45B pathway figure generated by IPA showing signaling components involved. Pink outline indicates molecules that are differentially expressed in *Egr3*-/-, compared with WT, mice following ECS. These include GADD45B and GADD45G. B. - C. Expression microarray results of GADD45B pathway gene expression in hippocampus from WT and *Egr3*-/- mice at baseline (No ECS) and 1 hour after ECS. B. *Gadd45b*, C. *Gadd45g*. D. -E. Quantitative RT-PCR validation of *Gadd45b* in D. The original male cohort and E. the replication female cohort. F. qRT-PCR validation of *Gadd45g* results in original male cohort. (n = 4-5 animals/group; *p<0.05,**p<0.01,***p<0.001, controlled for multiple comparisons). Statistical analyses for these, and all subsequent graphs, are shown in Table 3.

Expression microarray results showed that ECS causes a ≥2-fold increase in mRNA levels of both of these genes in WT mice that was not present in *Egr3*-/- mice (Fig. 3B- 3C). For both of these genes this resulted in a significantly lower level of expression following ECS in *Egr3*-/- mice than in WT controls. (Table 3 lists results of ANOVAs and post-hoc comparisons for all follow-up studies.)

**Table 3.** Statistics for microarray and qRT-PCR analyses.

| Gene | Two way ANOVA results | Post hoc comparison (Tukey's test) results<br>($p < 0.05$, **$p < 0.01$, ***$p < 0.001$, ****$p < 0.0001$) |
| --- | --- | --- |
| Gadd45b (M, microarray) | sig interaction of genotype & treatment $F(1, 12) = 18.39$, $p = 0.0011$ | WT vs WT ECS ***, WT ECS vs KO ECS *** |
| Gadd45b (M, qRT PCR) | sig interaction of genotype & treatment $F(1, 12) = 12.55$, $p = 0.0041$ | WT vs WT ECS ***, WT ECS vs KO ECS ** |
| Gadd45b (F, qRT PCR) | sig effect of genotype $F(1, 14) = 5.302$, $p = 0.0372$;<br>sig effect of treatment $F(1, 14) = 15.80$, $p = 0.0014$ | WT vs WT ECS **, WT ECS vs KO ECS * |
| Gadd45g (M, microarray) | sig interaction of genotype & treatment $F(1, 12) = 18.39$, $p = 0.0011$ | WT vs WT ECS ***, WT ECS vs KO ECS ** |
| Gadd45g (M, qRT PCR) | sig interaction of genotype & treatment $F(1, 12) = 9.875$, $p = 0.0085$ | WT vs WT ECS ***, WT ECS vs KO ECS * |
| Gadd45g (F, qRT PCR) | Not sig | N/A |
| Cdnk1a (M, microarray) | sig interaction of genotype & treatment $F(1, 12) = 7.316$, $p = 0.0191$ | WT vs WT ECS ****, KO vs KO ECS *, WT ECS vs KO ECS * |
| Cenpa (M, microarray) | sig interaction of genotype & treatment $F(1, 12) = 263.1$, $p < 0.0001$ | WT vs WT ECS ****, WT ECS vs KO ECS **** |
| Cenpa (M, qRT PCR) | sig interaction of genotype & treatment $F(1, 12) = 103.9$, $p < 0.0001$ | WT vs WT ECS ****, WT ECS vs KO ECS **** |
| Cenpa (F, qRT PCR) | sig interaction of genotype & treatment $F(1, 14) = 68.28$, $p < 0.0001$ | WT vs WT ECS ****, WT ECS vs KO ECS **** |
| Fos (M, microarray) | sig interaction of genotype & treatment $F(1, 12) = 10.92$, $p = 0.0063$ | WT vs WT ECS ****, KO vs KO ECS *, WT ECS vs KO ECS ** |
| Fos (M, qRT PCR) | sig interaction of genotype & treatment $F(1, 12) = 5.808$, $p = 0.0329$ | WT vs WT ECS **, WT ECS vs KO ECS * |
| Fos (F, qRT PCR) | sig effect of treatment $F(1, 14) = 8.392$, $p = 0.0117$ | WT vs WT ECS * |
| Fosb (M, microarray) | sig interaction of genotype & treatment $F(1, 12) = 63.79$, $p < 0.0001$ | WT vs WT ECS ****, WT ECS vs KO ECS **** |
| Fosb (M, qRT PCR) | sig interaction of genotype & treatment $F(1, 12) = 14.01$, $p = 0.0028$ | WT vs WT ECS ***, WT ECS vs KO ECS *** |
| Fosb (F, qRT PCR) | sig interaction of genotype & treatment F (1, 14) = 7.094, p = 0.0185 | WT vs WT ECS ***, WT ECS vs KO ECS ** |
| Junb(M, microarray) | sig interaction of genotype & treatment F (1, 12) = 18.22, p = 0.0011 | WT vs WT ECS ****, KO vs KO ECS *, WT ECS vs KO ECS *** |
| Junb(M, qRT PCR) | sig effect of treatment F (1, 12) = 6.247, p = 0.0280 | posthoc comparisons NS |
| Junb(F, qRT PCR) | sig effect of treatment F (1, 14) = 6.569, p = 0.0225; sig effect of genotype F (1, 14) = 4.608, p = 0.0498 | posthoc comparisons NS |
| Npas4(M, microarray) | sig interaction of genotype & treatment F (1, 12) = 24.11, p = 0.0004 | WT vs WT ECS ****, WT ECS vs KO ECS *** |
| Npas4(M, qRT PCR) | sig interaction of genotype & treatment F (1, 12) = 8.37, p = 0.0135 | WT vs WT ECS **, WT ECS vs KO ECS ** |
| Npas4 (F, qRT PCR) | sig effect of treatment F (1, 14) = 9.73 p=0.0075, sig effect of genotype F (1, 14) = 9.68, p = 0.0077 | WT vs WT ECS *, WT ECS vs KO ECS * |
| Nr4a2 (M, microarray) | sig interaction of genotype & treatment F (1, 12) = 15.90, p = 0.0018 | WT vs WT ECS ***, WT ECS vs KO ECS *** |
| Nr4a2 (M, qRT PCR) | sig interaction of genotype & treatment F (1, 12) = 7.050, p = 0.0210 | WT vs WT ECS ***, WT ECS vs KO ECS ** |
| Nr4a2 (F, qRT PCR) | sig interaction of genotype & treatment F (1, 14) = 5.230, p = 0.0383 | WT vs WT ECS * |
| Mef2c (M, microarray) | sig interaction of genotype & treatment F (1, 12) = 21.07, p = 0.0006 | WT vs WT ECS**, KO vs. WT ECS** |
| Mef2c (M, qRT PCR) | sig interaction of genotype & treatment F (1, 12) = 9.484, p = 0.0095 | posthoc comparisons NS |
| Mef2c (F, qRT PCR) | Not sig | N/A |
| Calb2 (M, microarray) | sig effect of genotype F (1, 12) = 48.99, p < 0.0001 | WT vs KO***, WT ECS vs. KO ECS** |
| Calb2 (M, qRT PCR) | sig effect of genotype F (1, 12) = 11, p = 0.0061 | posthoc comparisons NS |
| Calb2 (F, qRT PCR) | Not sig | N/A |

To validate these findings, we conducted qRT-PCR on the original mRNA samples that were used to perform the microarray analysis. Since the animals used in the microarray were all male, we replicated the study in female animals to determine if the gene expression changes were sex specific.

Expression of *Gadd45b* validated in the mRNA samples used for the microarray (Fig. 3D). In addition, in the female validation group *Gadd45b* showed the same pattern of a >2-fold increase in expression following ECS in WT mice that was absent in *Egr3-/-* mice (Fig. 3E). For *Gadd45g* the microarray findings were validated in the male mRNA samples (Fig. 3F) but did not show a significant difference between genotypes in the female replication cohort (data not shown, two-way ANOVA not significant, see Table 3). These results indicate that *Egr3* is required for activity-dependent upregulation of GADD45 family gene expression in the hippocampus of male mice and, in the case of *Gadd45b*, also in female animals.

### DNA damage response gene *Cenpa* upregulated 12-fold by ECS, which requires EGR3

Histone H3-like centromere protein A (CENPA), a protein essential for the initial stages of centromere assembly ^42^, was recently shown to be critical for efficient DNA repair *in vivo* ^43^. *Cenpa* showed the greatest degree of differential expression between *Egr3-/-* and WT mice of all the genes in the microarray dataset. This was due to a 12-fold upregulation of *Cenpa* in WT mice in response to ECS that was entirely absent in *Egr3-/-* mice (Fig. 4Ai, Table 3). This finding was validated by qRT-PCR in male mRNAs as well in the female replication cohort, in which ECS induced a 13-fold and 15-fold increase in *Cenpa* expression (respectively) in WT mice with no change in expression in *Egr3-/-* mice (Fig. 4Aii & 4Aiii, Table 3).

**Figure 4.**
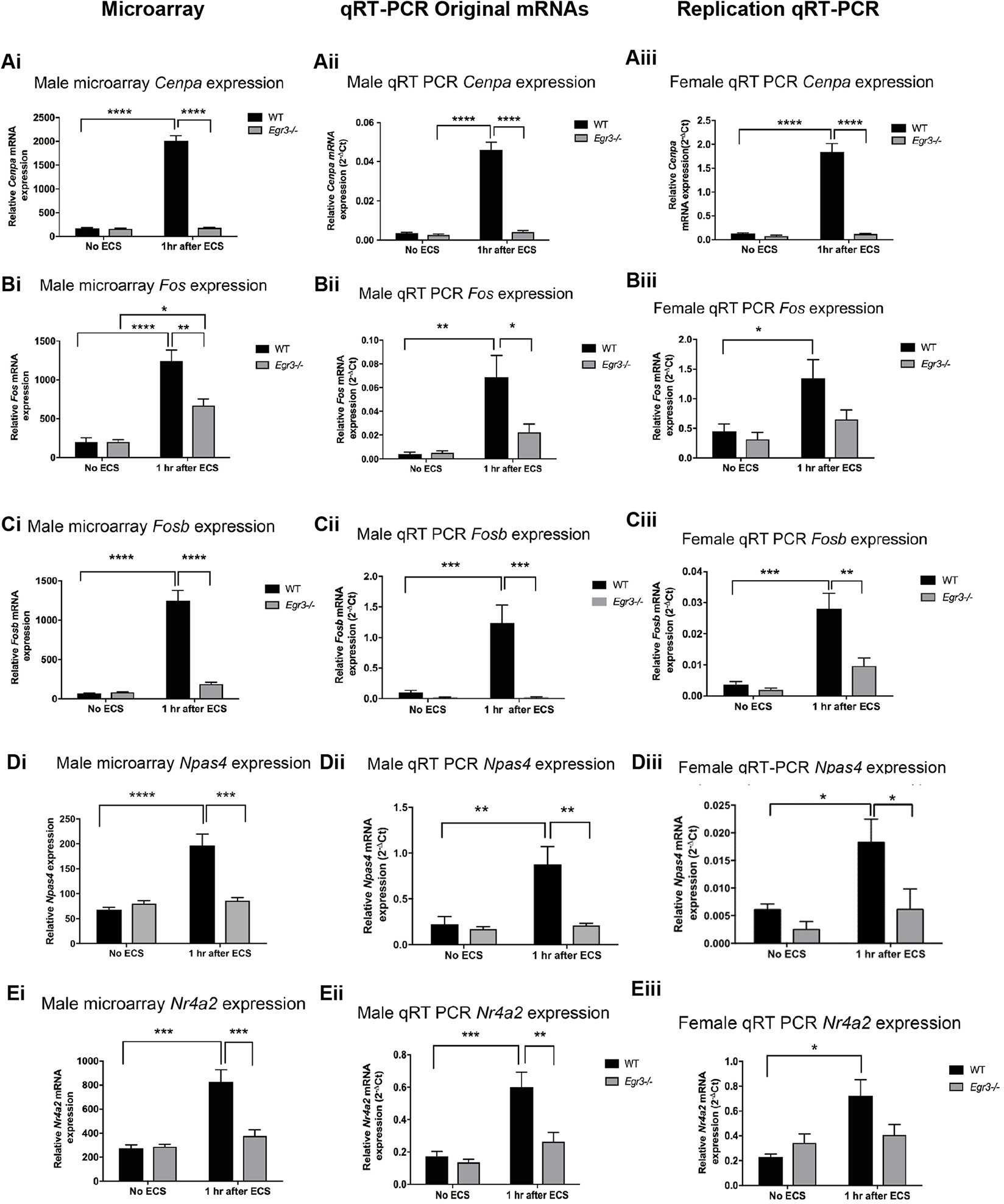
Numerous genes are differentially expressed in *Egr3*-/- mice compared with WT mice following ECS. Microarray analysis results (i) and follow-up qRT-PCR results performed in the original RNA samples used in the microarray (ii) and in a replication cohort of female mice (iii) for A. *Cenpa*, the most highly differentially expressed gene in the dataset, AP-1 family genes B. *Fos* and C. *Fosb*, and additional memory-related genes D. *Npas4* and E. *Nr4a2*, which is also involved in DNA repair. (n = 4-5 animals/group; *p<0.05,**p<0.01,***p<0.001, ****p<0.0001, controlled for multiple comparisons).

### ECS- induced gene expression of AP1 transcription factor components *Fos* and *Fosb* is *Egr3* dependent

The Activator protein 1 (AP1) transcription factor is a dimeric transcription factor whose subunits belong to four different families of DNA-binding proteins including the *Jun* family, *Fos* family, ATF/cyclic AMP-responsive element-binding (CREB) and the musculoaponeurotic fibrosarcoma (*Maf*) family ^44^. The AP-1 transcription factor components play critical roles in cancer^45^, immune system function^46^, neurite growth ^47^ and DNA repair ^48^. Results from the microarray showed that three AP-1 components *Fos, Fosb* and *Jun* showed ECS-induced expression WT mice that was either significantly lower, or absent, in *Egr3-/-* mice after ECS (Figs. 4Bi, 4Ci, Table 3). Both *Fos* and *Fosb* findings were validated in male RNA samples (Figs. 4Bii, 4Cii, Table 3) and replicated in the female cohort by qRT-PCR (Figs. 4Biii, 4Ciii, Table 3).

The initial microarray showed differential expression of *Jun* between *Egr3*-/- and WT mice. The follow-up qRT-PCR studies revealed a significant effect of genotype in the female replication cohort, but not in the male validation study. See Table 3.

### ECS-mediated induction of memory regulation genes *Npas4* and *Nr4a2* is *Egr3-* dependent

In addition to genes involved in DNA damage response, based on our IPA analysis results, the next category of genes that we decided to investigate further were those involved in nervous system function and regulation of behavior. Two genes, neuronal Per-Arnt-Sim domain 4 (*Npas4*) and nuclear receptor subfamily 4 member 2 (*Nr4a2*) particularly stood out due to their roles in memory and neurophysiologic processes. *Npas4* is essential for excitatory to inhibitory balance in the brain ^49^ and synapse maintenance ^50^ and plays important roles in working memory and cognitive flexibility ^51^. *Nr4a2* is required for both long-term memory and object recognition ^52^ and hippocampal neurogenesis ^53^ and was recently shown to be critical for DNA repair *in vivo*^*54*^.

In WT mice, ECS induced a 2.8 – 3-fold upregulation of both *Npas4* and *Nr4a2* expression that was absent in *Egr3*-/- mice (Figs. 4Di, 4Ei, Table 3). The results of the microarray were validated in the male RNAs and replicated in the female cohort by qRT-PCR for both genes (Figs. 4Dii – iii, 4Eii -iii, Table 3).

### Genes upregulated in *Egr3*-/-mice including *Mef2c* and *Calb2* are linked to schizophrenia

The majority of genes we chose for validation studies were upregulated in WT mice after ECS compared to *Egr3-/-* mice following ECS. However, a small number of genes showed increased expression in *Egr3-/-* mice compared with WT mice after ECS. We chose two genes from this group for validation studies based on their degree of fold-change induction, involvement in behavior and/or DNA damage response, and association with schizophrenia. The first, transcription factor myocyte enhancer factor 2c (*Mef2c*), is a key regulator of learning and memory *in vivo*^*55*^. The second, calbindin 2 (*Calb2*), encodes the protein calretinin, a critical regulator of long-term potentiation in the dentate gyrus^56^. Previous studies showed that enrichment of MEF2C motifs was found in sequences surrounding the top single nucleotide polymorphisms within schizophrenia risk loci ^57^ and increased levels of calretinin were reported in the dentate gyrus of schizophrenia and bipolar patients compared to controls^58^.

The *Mef2c* microarray results showed that ECS caused a decrease in *Mef2c* expression in WT mice which did not occur in *Egr3-/-* mice (Fig. 5A, Table 3). The opposing effect of ECS on *Mef2c* expression resulted in a significant interaction between ECS treatment and genotype in the two-way ANOVA in the microarray, a result that was validated by qRT-PCR in the male mRNAs (Fig. 5A – B, Table 3). In contrast, in females no significant changes in *Mef2c* expression were seen (Table 3).

**Figure 5.**
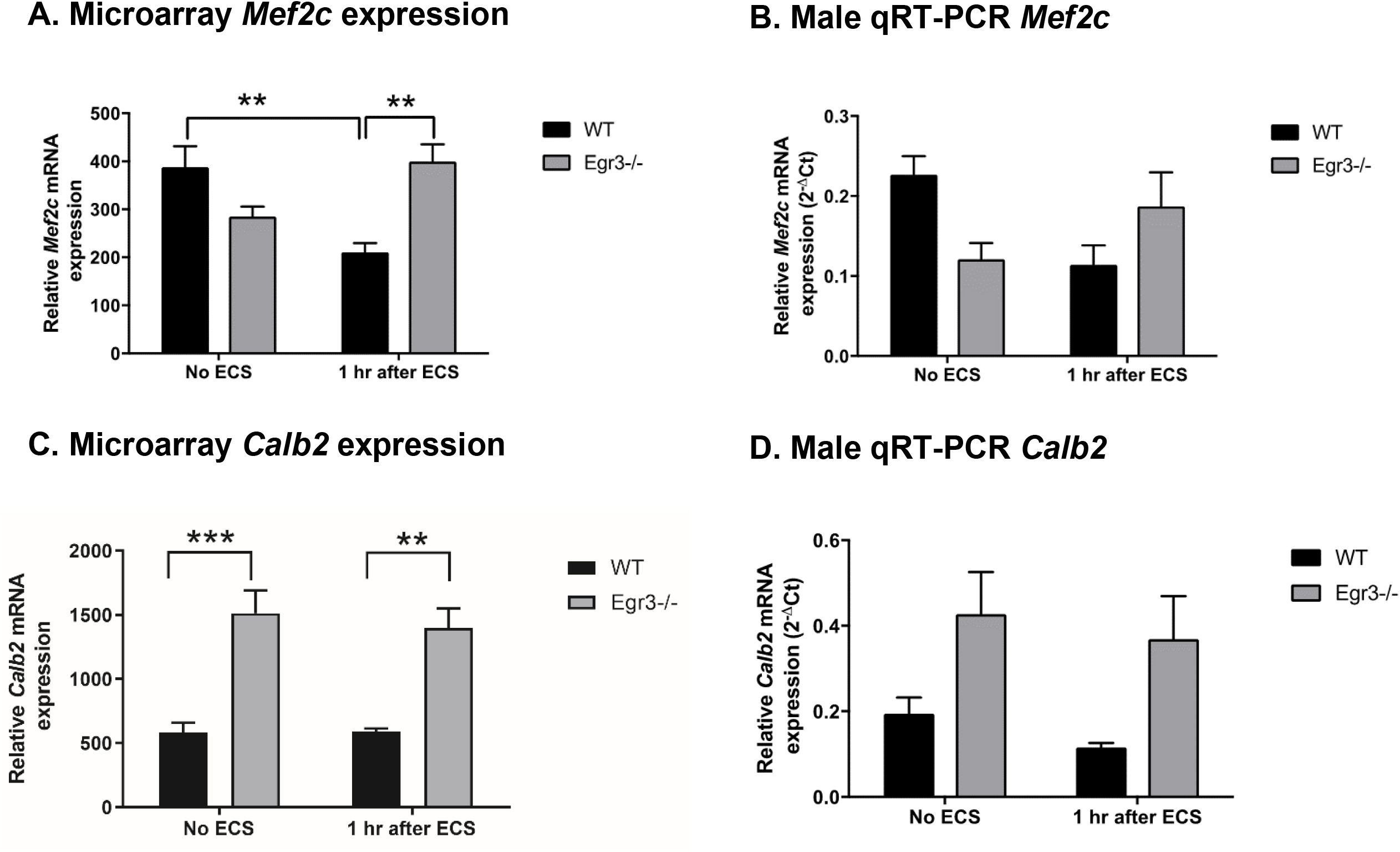
*Mef2c* and *Calb2* display unique patterns of regulation. Microarray analysis results (A and C) and follow-up qRT-PCR results in males (B and D) for schizophrenia associated gene *Mef2c* (A-B) and *Calb2* (C-D), genes overexpressed in schizophrenia. (n = 4 animals/group; **p<0.01, ***p<0.001, controlled for multiple comparisons).

*Calb2* expression levels showed a pattern that was rare in the dataset. Expression of *Calb2* is significantly reduced in *Egr3-/-* mice compared to WT mice at baseline and does not change in response to ECS in either genotype. This pattern, present in the microarray data and validation qRT-PCR in males (Fig. 5C – D, Table 3), was not significant in the female qRT-PCR results (Table 3).

## Discussion

Major advances in genetics and genomics over the last decade and a half have led to identification of hundreds of genes associated with risk for neuropsychiatric and neurodegenerative disorders. One method to identify mechanisms that unite these findings, and thus underlie illness etiology, is to identify the “master regulatory genes” that orchestrate expression of large numbers of these disease-associated genes. *EGR3* has been identified as such a master regulator in neuropsychiatric illnesses schizophrenia, bipolar disorder and, most recently, Alzheimer’s disease ^2-4^. These studies have relied on bioinformatics resources to identify the gene interaction relationships that led to these discoveries. However, few studies have validated the genes that require *Egr3* for their expression in the brain *in vivo*. Our findings reveal numerous genes that are dependent upon *Egr3* for their normal expression in response to neuronal activity in the mouse hippocampus. Numerous of these genes either map to schizophrenia GWAS risk loci or have been identified as differentially regulated in schizophrenia studies.

As an immediate early gene transcription factor *Egr3* is rapidly expressed in the brain in response to neuronal activity and, in turn, regulates the subsequent set of genes expressed in response to that activity. *Egr3* is thus poised to translate environmental stimuli into changes in gene expression that dictate the brain’s response to the outside world. Our studies in mice identified the critical roles of *Egr3* in stress-responsive behavior, memory, and synaptic plasticity ^13, 15^. Based on these findings, and the upstream signaling events that trigger *EGR3* expression, we hypothesized that dysfunction of *EGR3* would disrupt the brain’s resilient response to stress, resulting in neuropathology which, over time, may give rise to symptoms of neuropsychiatric illness ^13, 15, 16^.

Our hypothesis has subsequently been supported by studies showing both genetic association of *EGR3* with schizophrenia ^9, 59-61^, as well as decreased *EGR3* gene expression in brains of schizophrenia patients ^59, 62^ and fibroblasts isolated from schizophrenia patients ^63^. Recent *in silico* studies identified *EGR3* as a central gene in a network of transcription factors and microRNAs associated with schizophrenia risk ^2^, a master regulator of genes that are differentially regulated in bipolar disorder patients ^3^, and a critical regulator of DEGs involved in synaptic function in Alzheimer’s disease ^4^. In total, these findings suggest that altered *EGR3* activity, or disruption of proteins that function upstream or downstream of *EGR3*, may increase risk for neuropsychiatric disorders and play a role of development of neurodegenerative disease.

To identify *Egr3*-dependent genes in the brain, we used ECS to maximally induce immediate early gene expression in the hippocampi of WT and *Egr3*-/- mice and identified DEGs using a microarray-based approach. We found genes involved in regulation of behavior and DNA damage response pathways to be significantly overrepresented in our DEG list. Our results suggest that *Egr3* is necessary for ECS-dependent stimulation of a subset of genes involved in regulation of nervous system function including regulation of memory (*Npas4* ^*51*^, *Nr4a2* ^52^), neurogenesis and synaptic plasticity (*Gadd45b* ^*40*^, *Bdnf* ^64^), behavior (*Fos* ^65^, *Fosb* ^66^) and DNA damage response, particularly, DNA repair (*Cenpa* ^43^, GADD45 family proteins Gadd45b and *Gadd45g* ^37, 38^, *Fos* and *Fosb* that are part of the AP-1 transcription factor ^48^, and *Nr4a2* ^54^). We also report that two genes that show elevated expression in schizophrenia (*Calb2* ^*58*^, *Mef2c* ^57^) are upregulated in mice lacking *Egr3*. In total, our findings suggest that *Egr3* is critical for the normal activity-responsive expression of genes involved in brain function and the DNA damage response.

### The importance of DNA repair in neurons, findings of DNA damage regulating genes involved in behavior

In neurons, DNA damage can occur during normal cellular activity and in processes involving DNA replication, such as neurogenesis ^67^. Neurons are postmitotic, and typically cannot be replaced by new cells if DNA damage reaches critical levels. Therefore, neurons rely heavily on effective DNA repair mechanisms to maintain homeostasis ^68^. DNA damage is often associated with aging and disease pathology; however, two recent paradigm-shifting studies highlight the role of DNA damage in regulation of normal physiological function in neurons. The first study, by Suberbeille and colleagues^69^, showed that neuronal activity triggered by exploration of a novel environment can cause DNA damage in the form of DNA double-stranded breaks in the cortex and hippocampus of young adult WT mice. The second study, by Madabhushi and colleagues, demonstrated that *in vitro* stimulation of primary neurons induced DNA double-stranded breaks in the promoters of immediate early genes that was essential for their activity-dependent induction ^70^. They also showed that inhibiting non-homologous end joining, a DNA repair pathway, caused a sustained “switched on” state of gene expression perturbing the normal temporal dynamics of immediate early gene expression.

We found several genes from the Madabhushi study to be differentially expressed in our results. These included *Fos, Fosb, Nr4a2* and *Npas4*, which failed to be induced in *Egr3-/-* mice after ECS, and represented 4 of the 12 genes that showed upregulation in the Madabhushi study following etoposide treatment of neurons ^70^. Madabhushi and colleagues also reported that of these genes, *Fos, Fosb* and *Npas4* showed enrichment of DNA damage double strand breaks (γ-H2AX) in their promoters and were induced following neuronal activity *in vitro* ^70^.

We show that *Egr3* is necessary for the neuronal activity induced expression of these genes. Given the roles of these genes in regulation of neuronal function, impaired expression of these genes may contribute to the behavioral and cognitive deficits seen in *Egr3-/-* mice ^13, 71^. In addition to playing a role in behavior regulation, *Fos* and *Fosb* are members of the AP-1 transcription factor complex, a critical regulator of DNA repair genes ^48^. In line with these data, we also found that genes belonging to the GADD45 signaling pathway, a major DNA damage response pathway ^37, 38^, showed impaired induction in *Egr3-/-* mice following ECS. GADD45b was recently identified as an EGR3 dependent gene in prostate cancer, and EGR3 was shown to bind to the GADD45B promoter *in vivo* and upregulate expression of GADD45B *in vitro* ^72^.

In addition, other DNA damage response genes including *Cenpa* and *Nr4a2*, recently shown to play a role in DNA repair ^43, 54^ showed a similar lack of induction in the *Egr3-/-* mice after undergoing ECS. For several of these genes we saw particularly robust induction following ECS in wildtype mice including a 12-fold induction for *Cenpa* and a 15-fold induction for *Fos* that the *Egr3-/-* mice lacked. Given that *EGR3* is induced in response to DNA damaging stimuli *in vitro* ^*73*^, our findings suggest that lack of functional *Egr3* results in diminished activation of genes involved in DNA damage response. This dysfunction of *Egr3* may increase susceptibility to DNA damage, impacting normal physiological activation of genes and contributing to increased DNA damage observed in neuropsychiatric illness ^74-78^.

### Genes upregulated in *Egr3*-/- mice

While the majority of DEGs in our data failed to be induced by ECS in *Egr3-/-* mice, a small subset showed the opposite trend. Key among these genes were *Mef2c* and *Calb2*, which showed the most profound increase in *Egr3-/-* mice following ECS compared to WT mice. Prior studies show that both of these genes show increased expression in schizophrenia patients’ brains ^57, 58^. A previous study showed that deletion of *Mef2c* impairs hippocampal-dependent learning and memory *in vivo* ^55^. Also, *Mef2c* limits excessive synapse formation during activity-dependent synaptogenesis in the dentate gyrus ^55^. Mice lacking the *Calb2* encoded protein calretinin, show impaired hippocampal long-term potentiation (LTP) ^79^. Studies suggest that temporal and spatial regulation of MEF2 family members ^80^ and *Calb2* ^81^ is essential for normal brain development.

Both timing and level of immediate early gene expression are critical features of their function. Either insufficient expression, or persistent overexpression, of immediate early genes, or the factors that regulate them, can negatively affect learning ^82^, or cause anxiety-like behavior, respectively ^83^. Our findings indicate that *Egr3* may influence the temporal regulation of these genes, where a lack of *Egr3* may lead to a perpetual “switched on” state of gene expression for genes such as *Mef2c* and *Calb2* that is not observed in WT mice, and may negatively impact normal brain function.

In summary, we report activity-dependent gene expression changes in the hippocampus of *Egr3-/-* mice, previously shown to exhibit schizophrenia-like behavioral abnormalities and memory deficits ^13, 15^. We have validated numerous genes that were differentially expressed in the microarray data using qRT-PCR in the original male RNAs and a replication cohort of female mice, demonstrating that many of these effects are independent of sex while others appear sex-dependent. These genes are involved in behavior and DNA damage response, with a subset of these playing dual roles in neuronal function and DNA repair including *Gadd45b, Nr4a2* and *Bdnf*. Further studies are needed to define the role of EGR3 in regulating DNA response genes necessary to repair DNA double strand breaks induced by neuronal activity. In conclusion, our studies demonstrate that EGR3 is a critical dual regulator of behavior and DNA damage response genes, and further define its role in brain function and neuropsychiatric and neurodegenerative illnesses characterized by cognitive dysfunction. The identification of EGR3-dependent genes in the mouse hippocampus may help to explain finding indicating that EGR3 may be a master regulator of genes differentially expressed in neuropsychiatric illnesses ranging from schizophrenia and bipolar disorder to Alzheimer’s disease ^2-4^.

## Acknowledgements

The authors gratefully acknowledge the assistance of past and present members of the Gallitano Lab Aneri Mehta, Annika Ozols, and Xiuli Zhao, PhD. Research reported in this publication was supported by the National Institute of Mental Health of the National Institutes of Health (NIH) under Award Numbers R01MH097803 and R21MH113154 (to ALG), and R01MH107487 and R01MH121102 (to RRM), and by the National Institute on Aging of NIH under Award Number R01AG057598 (to RRM). The content is solely the responsibility of the authors and does not necessarily represent the official views of the NIH.

