## Supplemental Materials for "Identification of activity-induced *Egr3*-dependent genes reveals genes associated with DNA damage response and schizophrenia"

#### Supplemental Figures

#### Figure S1

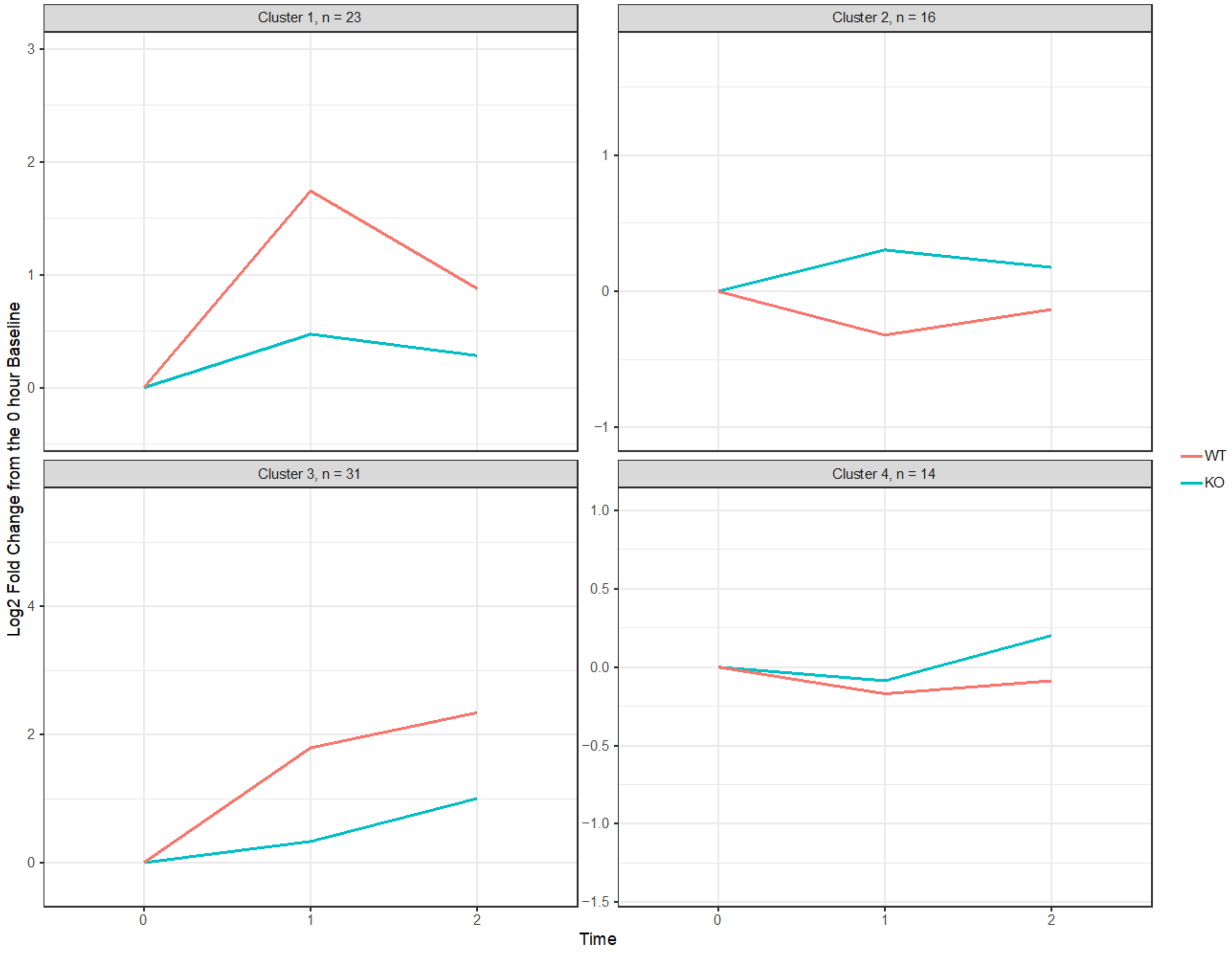


**Figure S1. Differentially expressed genes fall into four clusters.**
The four clusters of DEGs generated by the k-mean clustering analysis are visualized here to represent the pattern of gene expression changes across the different timepoints relative to the baseline (timepoint = 0). The number of clusters (k) was determined using the “elbow” method. The n in the headings of each panel represents the number of genes in each cluster. The transparent lines show the log2 fold change relative to the baseline for each gene, whereas the highlighted lines show the average of these patterns per group. The colors denote the two groups (KO: *Egr3*-/- and WT).

**Figure S2**

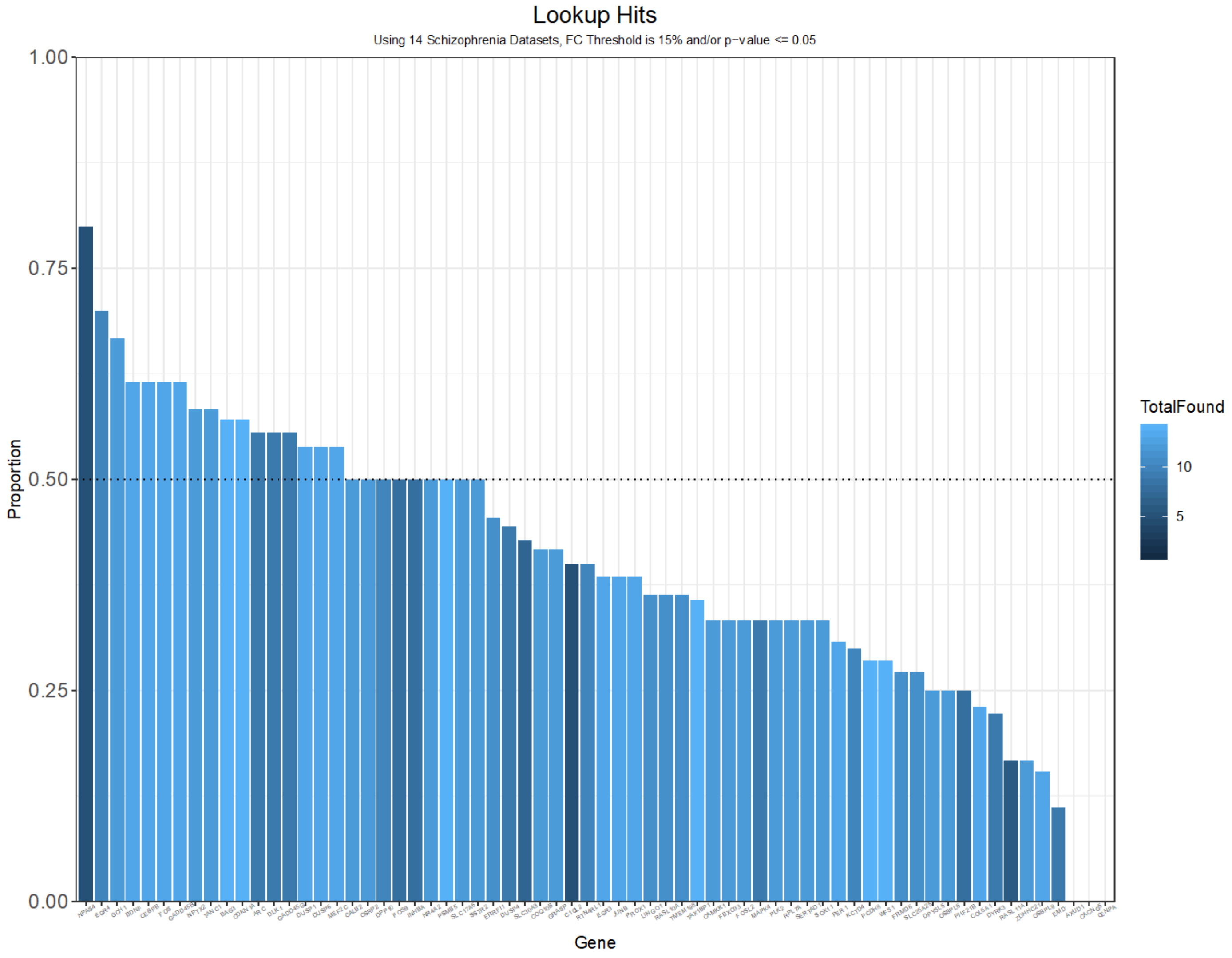


**Figure S2. Proportion of schizophrenia studies in which each DEG is identified**
Fourteen published gene expression studies in schizophrenia were queried for the 71 DEGs in *Egr3*-/- compared with WT mouse hippocampus 1 hr following ECS. Histogram plot showing the proportion of the 14 studies in which each gene was found to be differentially expressed in schizophrenia patient samples compared with controls (fold change is at least 15% and/or p-value <= 0.05). The gene NPAS4 was the most commonly identified DEG, appearing in 80% of the 14 schizophrenia datasets.

### Table S1. Primers for qRT-PCR

| Gene | Forward | Reverse | Reference |
|---|---|---|---|
| Gadd45b | GTT CTG CTG CGA CAA TGA CA | TTG GCT TTT CCA GGA ATC TG | Ma et al., Science 2009, Feb 20;323(5917):1074-7, PMID: 19119186. |
| Gadd45g | TCG CAC AAT GAC TCT GGA AG | CAG GGT CCA CAT TCA GGA CT | Ozawa et al., J. Toxicological Sci. 2011, Vol 36, No. 5, 613-623, PMID: 22008536.<br>Gavin et al., Epigenomics 2015, 7(4):567-569, PMID: 26111030. |
| Cdkn1a | GCA AAG TGT GCC GTT GTC | AGA CCA ATC TGC GCT TGG | Yamaguchi et al., Genes Dev. 2010, Mar 1;24(5):455-69, PMID: 20194438. |
| Cenpa | TTG GCC CTT CAG GAG GCA GCA | AAG CGT GAC CCG ACC AGC AT | McGregor et al., Cell Cycle 2014;13(5):739-48, PMID: 24362315. |
| Nr4a2 | ACA CAC ACA CCT TAA TGG GAC CCT | CAT GCC ACC CAC GCA ACA TTT AGT | Lundequist et al., Mol Immunol. 2011 Sep;48(15-16):1753-61, PMID: 21621845. |
| Fosb | AGG CAG AGC TGG AGT CGG AGA T | GCC GAG GAC TTG AAC TTC ACT CG | Alibhai et al., Brain Res. 2007, Vol 1143:22-33, PMID: 17324382. |
| Fos | GAA CGG AAT AAG ATG GCT GC | TTG ATC TGT CTC CGC TTG G | Nott et al., Nat Neurosci. 2016, Nov;19(11):1497-1505, PMID: 27428650. |
| Sertad1 | CTC CCT CTT CGT TCT GAT TGG | AGA GGG CTT CCA TCG TCT | Harouz et al., EMBO J. 2014, Nov 18;33(22):2606-22, PMID: 25216677. |
| Junb | AGG CTA GCT TCA GAG ATG CG | ATC CCT ATC GGG GTC TCA AG | Zhang et al., Plos One 2014, Dec 3;9(12):e114071, PMID: 25470242. |
| Npas4 | CTG CAT CTA CAC TCG CAA GG | GCC ACA ATG TCT TCA AGC TCT | Ramamoorthi et al., Science 2011, Dec 23; 334(6063):1669-1675, PMID: 22194569. |
| Pgk1 | TGT TAG CGC AAG ATT CAG CTA GTG | CAG ACA AAT CCT GAT GCA GTA AAG AC | Maple et al., ACSChem Neurosci. 2015, 6 (7), pp 1137–1142, PMID: 25857407. |
